# Small Molecule Binding to EDS1/PAD4 in LRR-RP-Mediated Pattern-Triggered Immunity

**DOI:** 10.64898/2025.12.10.692544

**Authors:** Judith Fliegmann, Denis Janocha, Chenlei Hua, Edda von Roepenack-Lahaye, Mark Stahl, Federica Locci, Jane E. Parker, Thorsten Nürnberger, Lisha Zhang

**Affiliations:** Department of Plant Biochemistry, Centre of Plant Molecular Biology (ZMBP), Eberhard-Karls-University of Tübingen, Tübingen, Germany; Analytics Unit, Centre of Plant Molecular Biology (ZMBP), Eberhard-Karls-University of Tübingen, Tübingen, Germany; Department of Plant-Microbe Interactions, Max Planck Institute for Plant Breeding Research, Cologne, Germany

## Abstract

Pattern-triggered immunity (PTI) is a key defense mechanism initiated when pathogen-associated molecular patterns are recognized by receptors, including leucine-rich repeat receptor proteins (LRR-RPs)^1,2^. EDS1 and PAD4 are crucial for LRR-RP-mediated PTI activation^3^. Small molecule (SM) phosphoribosyl-AMP/ADP, generated by TIR-domain NADase activity, promote the recruitment of helper NLRs into EDS1-PAD4 complex^4,5^. Here, we show that BAK1 recruitment to LRR-RP complexes is unaffected in *eds1-2* or *pad4-1* mutants, but is abolished in the *sobir1-12* mutant, highlighting the essential role of SOBIR1 in surface immune receptor complex formation. Furthermore, SM-binding to EDS1 and PAD4 is critical for LRR-RP-mediated PTI, as SM-binding-deficient mutants fail to restore immune responses. De novo SM synthesis is required for late PTI responses, including *PR1* expression and bacterial resistance, but not for early responses such as ROS and ethylene production. The TNL SADR1 functions only in late PTI responses, suggesting distinct roles of multiple TIR-domain proteins in PTI signaling.

---

Plants defend against microbial infection through two major classes of immune receptors: plasma membrane–localized pattern recognition receptors (PRRs) that activate PAMP-triggered immunity (PTI), and intracellular NLR receptors that mediate effector-triggered immunity (ETI)^6,7^. PTI and ETI act in concert to provide robust resistance against pathogen attack^2,3,8-10^.

In *Arabidopsis thaliana*, cell-surface leucine-rich repeat (LRR) receptor kinases (LRR-RKs) and LRR receptor proteins (LRR-RPs) primarily recognize proteinaceous PAMPs^1,11^. For example, the LRR-RKs FLS2 and EFR detect flg22 and elf18, whereas the LRR-RPs RLP23, RLP30, RLP32, and RLP42 recognize nlp20, small cysteine-rich proteins (SCPs), translation initiation factor 1 (IF1), and pg13, respectively^12-18^. Although LRR-RPs share ectodomain features with LRR-RKs, they lack an intracellular kinase domain and instead constitutively associate with the adaptor LRR-RK SUPPRESSOR OF BRASSINOSTEROID INSENSITIVE 1 (SOBIR1). Upon PAMP perception, both LRR-RKs and LRR-RPs recruit SERK-family co-receptors, including BAK1, to initiate intracellular signaling through receptor-like cytoplasmic kinases (RLCKs)^19^. LRR-RP– mediated signaling often differs from LRR-RK pathways in both kinetics and amplitude, with RLCKs PBL30 and PBL31 acting as key positive regulators of LRR-RP–triggered PTI^3,20^.

The lipase-like protein ENHANCED DISEASE SUSCEPTIBILITY 1 (EDS1) functions with either PHYTOALEXIN DEFICIENT 4 (PAD4) or SENESCENCE-ASSOCIATED GENE 101 (SAG101) to control distinct branches of NLR-mediated immunity^21^. During ETI, TIR-domain NLRs (TNLs) act as NADases that generate small signaling molecules (SMs)^5,22-25^. These SMs bind selectively to EDS1–PAD4 or EDS1–SAG101 heterodimers, inducing conformational changes that activate specific helper NLRs (hNLRs). For example, ADPr-ATP and di-ADPR activate the EDS1– SAG101–NRG1 module^22,26,27^, whereas pRib-AMP/ADP activates the EDS1–PAD4–ADR1 module^4,5^.

Recent work has broadened this framework, revealing that TIR-dependent signaling components—EDS1, PAD4, and ADR1—not only mediate ETI but also contribute centrally to early PTI activation^3^. Mutants defective in TIR signaling show attenuated PTI outputs and reduced pathogen resistance, underscoring an unexpected mechanistic overlap between PTI and ETI pathways^9,28^.

Here, we show that binding of TIR-derived small molecules to the EDS1–PAD4 complex is crucial not only for the onset of early PTI responses—such as rapid ROS generation and ethylene biosynthesis—but also for sustained defense outputs, including transcriptional activation of immune marker gene expression and disease resistance priming. Using mutants defective in SM binding, we find that disruption of this interaction compromises both immediate signaling events and later stages of PTI reinforcement, demonstrating that SM engagement acts as a central biochemical switch in LRR-RP–mediated immunity. Our findings further suggest that at least two distinct TIR-domain proteins contribute to PTI initiated by LRR-RP-type PRRs. We propose that one of these TIR proteins functions as a constitutive or low-level producer of SMs, thereby maintaining a basal pool of SMs that enable the EDS1–PAD4 complex to recruit and activate ADR1-family helper NLRs upon PRR stimulation. A second TIR protein, SADR1^28^, may act in a stimulus-dependent manner, amplifying SM production following PAMP perception. Together, these activities would establish a tunable signaling circuit in which constitutive and inducible TIR outputs converge on EDS1–PAD4 to potentiate helper NLR activation, thereby enabling robust and sustained PTI signaling downstream of LRR-RPs.

Perception of PAMPs by LRR-RPs initiates PTI through recruitment of the co-receptor BAK1 into cell surface immune receptor complexes. To determine whether the diminished PTI responses observed in *eds1-2* and *pad4-1* mutants^3^ arise from impaired BAK1 recruitment, we generated stable transgenic *Arabidopsis* lines in the Col-0, *eds1-2, pad4-1*, and *sobir1-12* backgrounds expressing either *p35S:RLP23:GFP or p35S:RLP42:GFP*. Fully expanded leaves from T2-generation plants were infiltrated with their cognate elicitors (nlp20 or pg13) or with water as a mock control. Five minutes after infiltration, GFP-tagged receptor complexes were immunoprecipitated using GFP affinity beads, and the presence of endogenous BAK1 in the complexes was assessed by immunoblotting.

Ligand-induced association of BAK1 with both RLP23–GFP and RLP42–GFP was robust in Col-0, *eds1-2*, and *pad4-1*, demonstrating that BAK1 recruitment is unaffected by loss of EDS1 or PAD4 (Fig. 1). In striking contrast, BAK1 recruitment was completely abolished in the *sobir1-12* background. These results establish that the LRR-RP–SOBIR1 complex is required for BAK1 recruitment and for initiation of downstream immune signaling, whereas EDS1 and PAD4 are dispensable for this early PTI event. This outcome was unexpected because previous biochemical analyses suggested that RLP23–BAK1 complexes could assemble *in vitro* upon nlp20 addition even in the absence of SOBIR1^15^. In contrast, our current *in vivo* genetic evidence indicates that functional SOBIR1 is essential for ligand-dependent recruitment of BAK1 into LRR-RP–SOBIR1 complexes.

**Fig. 1.**
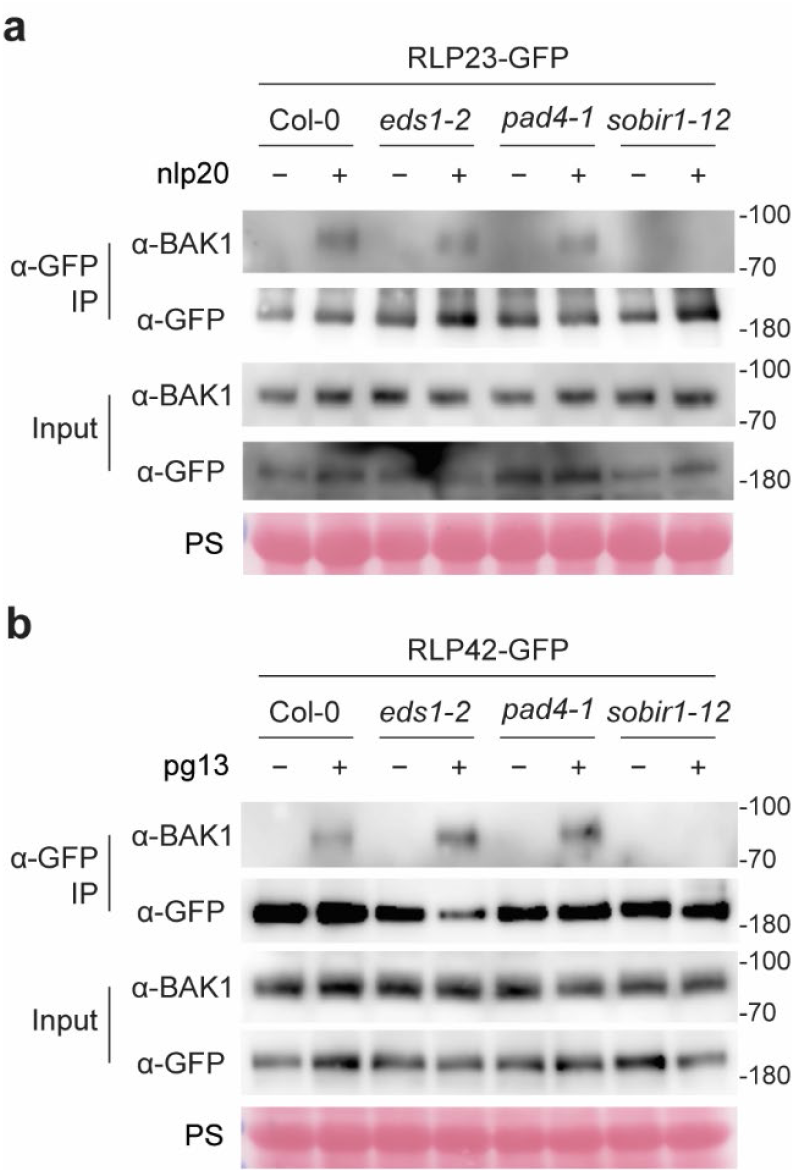
Recruitment of BAK1 into a complex with RLP23 and RLP42 requires SOBIR1, independent of EDS1 or PAD4, in Arabidopsis. Leaves from stable transgenic lines of Col-0, *eds1-2, pad4-1*, and *sobir1-12* overexpressing RLP23-GFP (**a**) and RLP42-GFP (**b**) were infiltrated with water (−) or 1 µM elicitor (+) for 5 min and subsequently harvested. Proteins extracted from these leaves were subjected to co-immunoprecipitation using GFP-trap agarose beads, followed by immunoblotting with anti-GFP or anti-BAK1 antibodies. Each experiment was replicated at least three times, yielding consistent results. Ponceau S (PS) staining was used as a loading control.

Earlier work demonstrated that wild-type EDS1—but not an SM-binding–deficient variant, EDS1^R493A^—restores nlp20-triggered ethylene production in *eds1-2*^3^, suggesting a critical role for SM binding in LRR-RP–triggered PTI. To extend this analysis, we complemented *eds1-2* with either wild-type EDS1 or EDS1^R493A 29^ and assessed responses to multiple elicitors. The *eds1-2* mutant showed markedly reduced reactive oxygen species (ROS) burst and ethylene production in response to the LRR-RP–recognized peptides nlp20, IF1, and pg13, but not to flg22, which is perceived by the LRR-RK FLS2 (Fig. 2a,b and Supplementary Fig. 1). Complementation with wild-type EDS1 restored LRR-RP–mediated responses, whereas EDS1^R493A^ failed to do so. Likewise, ligand-induced expression of the immune marker *PATHOGENESIS-RELATED 1* (*PR1*) was diminished in both *eds1-2* and *EDS1*^*R493A*^*/eds1-2* plants (Supplementary Fig. 2). In pathogen resistance assays, both nlp20 and pg23 (a 23–amino acid immunogenic core fragment of fungal polygalacturonases, including pg13)^14^ conferred resistance to *Pseudomonas syringae* pv. *tomato* (*Pst*) DC3000 in Col-0 and *eds1-2* complemented with wild-type EDS1, but not in *eds1-2* or *EDS1*^*R493A*^*/eds1-2* plants (Fig. 2c and Supplementary Fig. 3). Flg22-induced resistance remained unchanged across genotypes (Fig. 2c).

**Fig. 2.**
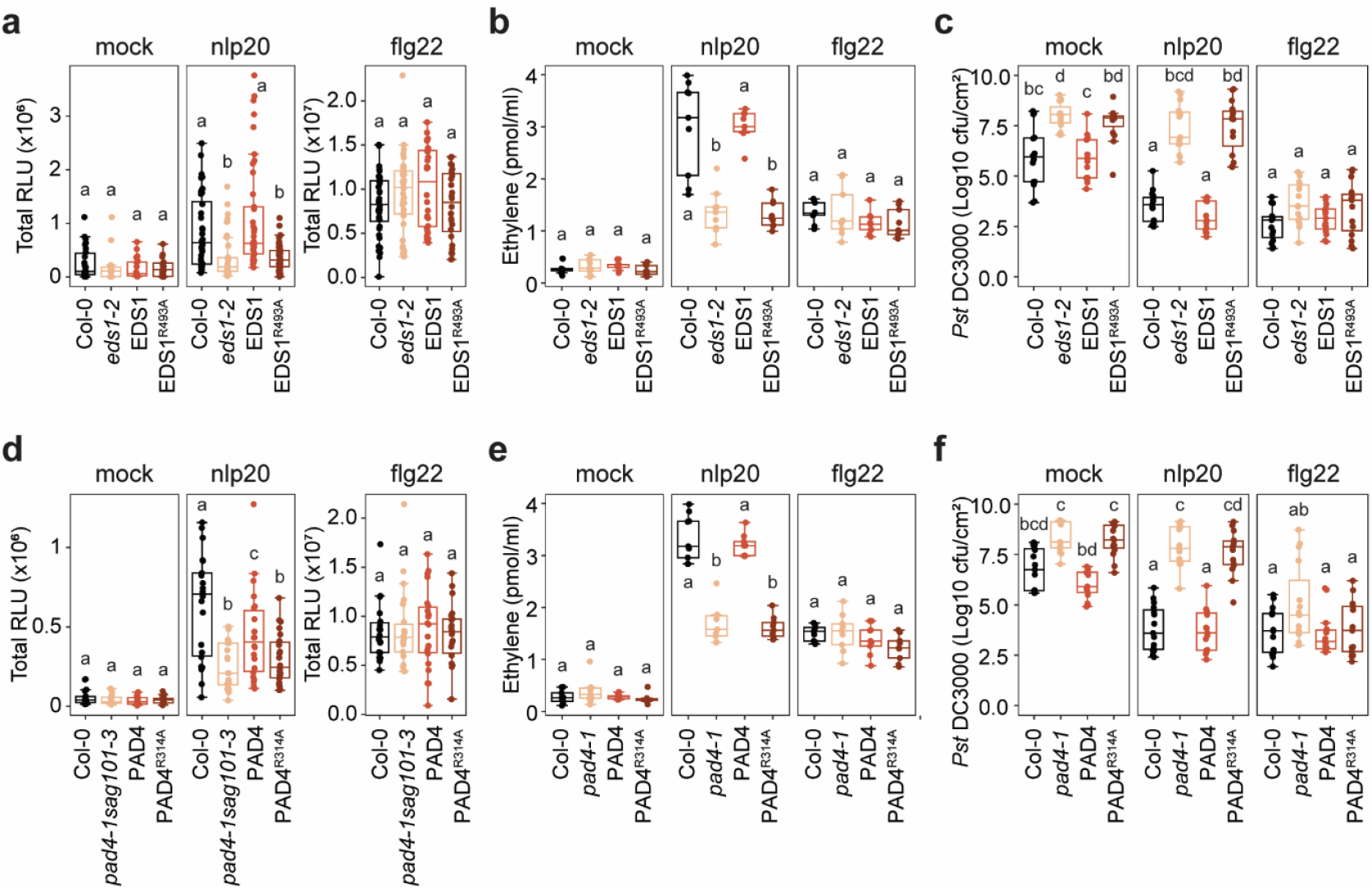
SM binding to EDS1 (**a-c**) and PAD4 (**d-f**) is essential for LRR-RP-mediated PTI. **a**,**d**, Total reactive oxygen species (ROS) production over 40 minutes in the indicated plants following treatment with water (mock) or 1 µM elicitor. Data represent four independent experiments. **b**,**e**, Elicitor-induced ethylene production after 4 h treatment with water (mock) or 1 µM elicitor. N = 9 from three independent experiments. **c**,**f**, Elicitor-induced defense against *Pst* DC3000. Leaves were treated with water (mock) or 1 µM elicitor 24 hours prior to *Pst* DC3000 infection. Bacterial titers were quantified in leaf extracts at 3 days post-inoculation (dpi). n = 15 biological replicates (each comprising two leaf discs). Letters indicate statistically significant differences (for **a**,**b**,**d**,**e**, among plant genotypes’ responses to each elicitor), determined by the Kruskal-Wallis test followed by post-hoc Dunn’s test (P < 0.01).

PAD4 residue R314 is likewise essential for SM binding. Using *pad4-1* and *pad4-1 sag101-3* lines complemented with either wild-type PAD4 or PAD4^R314A 30^, we observed that only wild-type PAD4 restored ROS and ethylene responses to nlp20, IF1, and pg13 (Fig. 2d,e and Supplementary Fig. 4). Nlp20-induced *PR1* expression and nlp20- and pg23-mediated resistance to *Pst* DC3000 were also recovered only in lines expressing wild-type PAD4 (Fig. 2f and Supplementary Fig. 2,3). Flg22-triggered responses were unaffected by PAD4 mutation. These results demonstrate that SM binding to both EDS1 and PAD4 is indispensable for LRR-RP–mediated PTI.

Interestingly, nlp20-triggered MAPK activation was impaired in *eds1-2* and was rescued by wild-type EDS1 but not EDS1^R493A^ (Supplementary Fig. 5a). In contrast, MAPK activation remained intact in *pad4-1* and *pad4-1 sag101-3* regardless of the complementing construct (Supplementary Fig. 5b,c). Protein levels of MPK3, MPK4, and MPK6 were comparable across Col-0, *eds1-2*, and *pad4-1* (Supplementary Fig. 6), indicating that SM binding to EDS1 is specifically required for MAPK activation in LRR-RP signaling.

TIR-domain proteins generate immune-stimulatory small molecules such as pRib-AMP/ADP upon effector perception through NADase activity, a process inhibited by excess nicotinamide (NAM)^28,31,32^. To determine whether such molecules are synthesized during PTI, we pretreated or co-treated plants with 50 mM NAM before elicitor application. Early immune responses—including ROS burst, MAPK activation, and ethylene production induced by nlp20, pg13, or flg22—were unaffected by either 16-hour NAM pretreatment or NAM co-infiltration (Fig. 3a-c and Supplementary Fig. 7). In contrast, nlp20-induced *PR1* expression at 24 hpi was significantly reduced by NAM (Fig. 3d), and NAM abolished nlp20-induced resistance to *Pst* DC3000 (Fig. 3e) as reported earlier^28^. These findings indicate that de novo synthesis of SMs is dispensable for early PTI responses but is required for late immune outputs.

**Fig. 3.**
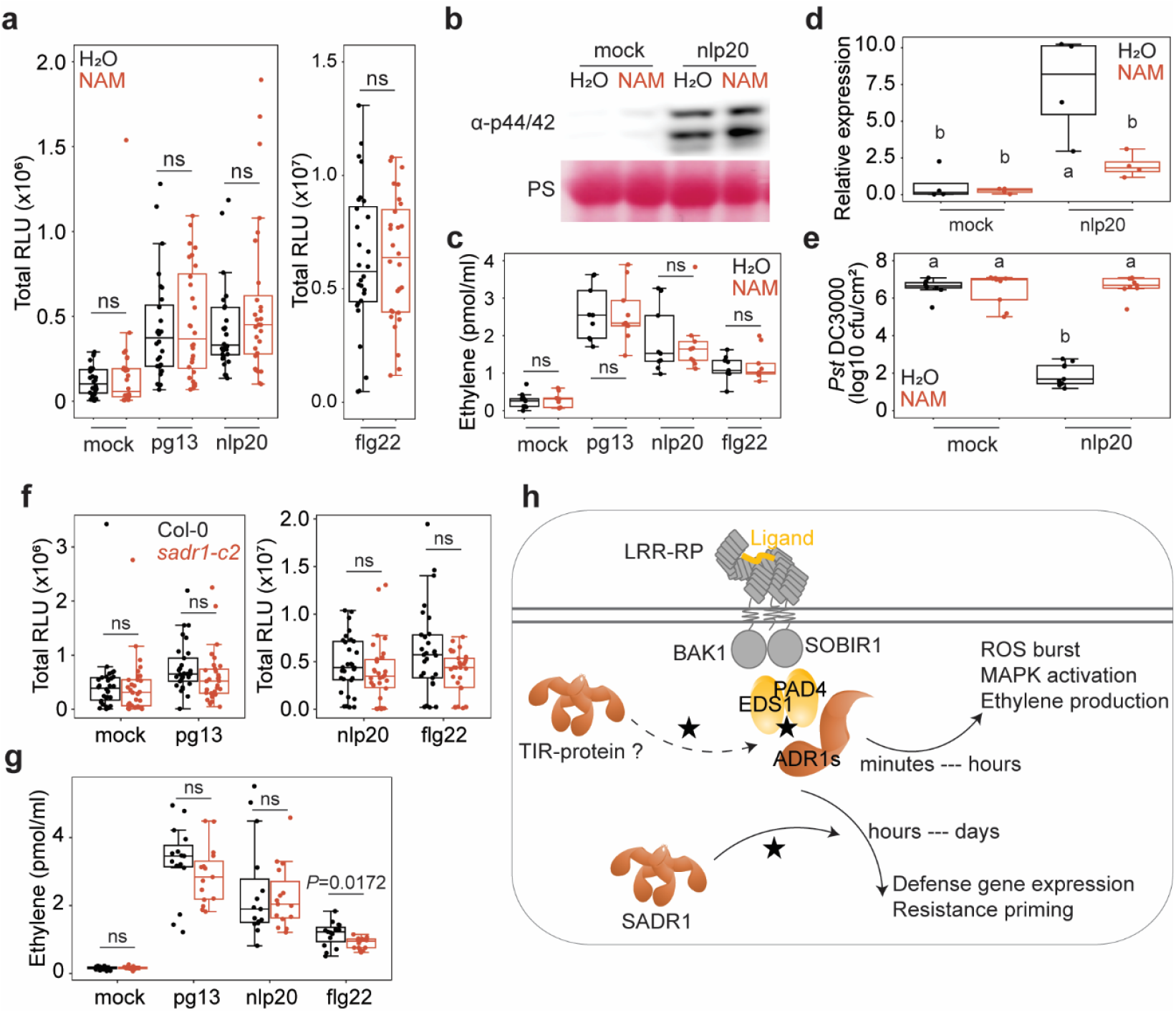
Differential regulation of TNL or TIR-domain proteins in PTI. **a-e**, De novo SM biosynthesis is required for late, but not early, PTI. **a**, Total ROS production over 60 minutes in Col-0 pre-treated with water or 50 mM NAM 16 hours prior to mock or elicitor treatment (N = 28 from 4 independent experiments). **b**, MAPK activation in Col-0 treated with mock or 1 µM nlp20 ± 50 mM NAM, detected by immunoblot using an anti-p44/42 MAP kinase antibody. Ponceau S (PS) staining was used as a loading control. Experiments were performed three times with consistent results. **c**, Ethylene accumulation after 5 hours of treatment with mock or 1 µM elicitor ± 50 mM NAM in Col-0 (N = 9 from 3 independent experiments). **d**, PR1 expression 24 hours post-treatment with water (mock) or 1 µM nlp20 ± 50 mM NAM (N = 4 biological replicates). **e**, *Pst* DC3000 resistance in Col-0 pre-treated with mock or 1 µM nlp20 ± 50 mM NAM. Bacterial titers were quantified in leaf extracts at 3 days post-inoculation (dpi). N = 9 biological replicates (each comprising two leaf discs). **f**,**g**, TNL SADR1 is not required for early PTI. **f**, Total ROS production over 60 minutes in Col-0 and *sadr1-c2* mutant plants, following treatment with water (mock) or 1 µM elicitor. Data represent five independent experiments (N = 30). **g**, Elicitor-induced ethylene production in Col-0 and *sadr1-c2* mutant plants. N = 15 from five independent experiments. **h**, Model of two functional layers of TNL or TIR-domain proteins contributing to LRR-RP-mediated PTI: i) a basal layer of TIR domain proteins (dashed line) produces low levels of SM (black star), facilitating the preassembly of the SOBIR1-EDS1-PAD4-ADR1 complex for early PTI responses (e.g., ROS burst, MAPK activation, and ethylene production); ii) a second layer involves inducible TIR domain proteins or TNLs, such as SADR1, which are activated downstream of early PTI signaling and are required for late PTI outputs (e.g., defense gene expression and resistance priming). Letters indicate statistically significant differences (ANOVA with post-hoc Tukey [**d**,**e**] or two-tailed Student’s t-test [**a**,**c**,**f**,**g**], P < 0.05; ns, not significant).

Given the requirement for de novo SM synthesis in late PTI, we further investigated whether TNL proteins contribute to this process. SADR1, a TNL known to be involved in nlp20-induced immunity^28^, was examined for its role in early PTI signaling. The *sadr1-c2* mutant exhibited normal reactive oxygen species (ROS) burst and ethylene production in response to the pathogen-derived elicitors pg13, nlp20, and flg22, showing similar levels of activity to the Col-0 wild-type control (Fig. 3f,g). These findings suggest that SADR1 is dispensable for the early-phase immune responses, which include ROS production and ethylene synthesis. However, SADR1 is crucial for the activation of late-stage immune responses, such as the expression of *PR1* and disease resistance priming^28^, emphasizing its specialized role in the later stages of pattern-triggered immunity (Fig. 3h).These findings also suggest the existence of other TNL or TIR-domain proteins involved in early PTI processes that are independent of SADR1.

In our study, we demonstrate that endogenous SOBIR1, rather than EDS1 or PAD4, plays a crucial role in recruiting BAK1 to the LRR-RPs RLP23 and RLP42 upon sensing their ligands nlp20 and pg13 (Fig. 1). Previous *in vitro* analyses using size exclusion chromatography showed that the ectodomains of RLP23 and BAK1 can form a ligand-induced complex in the presence of nlp20, indicating a direct extracellular interaction between these proteins ^15^. Similarly, structural work on the *Nicotiana benthamiana* LRR-RP RXEG1 revealed a ternary complex with its ligand XEG1 and BAK1 assembled through ectodomain contacts^33^. Notably, SOBIR1 was absent in both *in vitro* systems, suggesting that LRR-RP–BAK1 associations can form independently of SOBIR1 under these conditions. However, our *in vivo* data demonstrate that SOBIR1 is required to facilitate or stabilize BAK1 recruitment to LRR-RP complexes following ligand perception. Together, these findings support a model in which SOBIR1 functions as a scaffold or co-receptor that promotes formation of an active LRR-RP–SOBIR1–BAK1 signaling complex at the plasma membrane. To resolve the architecture and dynamics of this assembly, future structural studies incorporating a tetrapartite complex of the LRR-RP, ligand, SOBIR1, and BAK1 will be essential.

PTI is initiated within minutes of pathogen detection, resulting in rapid cellular reprogramming and extensive transcriptional changes during the first hour after elicitation^34^. In contrast, ETI generally becomes active several hours after pathogen encounter. During ETI, TNLs act as NADases that generate SMs such as pRib-AMP/ADP, which bind to the EDS1–PAD4 heterodimer to recruit the helper NLR ADR1^4,5^. This EDS1–PAD4–ADR1 signaling module has been classically associated with amplifying immune responses during ETI. We and others previously established that this module also contributes to PTI^3,9^. Here, we further show that SM binding to both EDS1 and PAD4 is required to activate downstream immune responses. Point mutations that disrupt SM binding in EDS1 (R493A) and PAD4 (R314A) result in reduced ROS production, impaired ethylene accumulation, strongly diminished PR1 expression, and attenuated resistance to *Pst* DC3000, underscoring the importance of SM binding for both early and late PTI responses (Fig. 2).

Our observation that the EDS1–PAD4–ADR1 complex associates with SOBIR1 in a ligand-independent manner further suggests the existence of a pre-formed immune complex composed of the LRR-RP, SOBIR1, and the EDS1–PAD4–ADR1 module in unchallenged plants^3^. This model is supported by our nicotinamide (NAM) inhibition experiments. Blocking TNL NADase activity with NAM selectively impairs late PTI outputs, including PR1 expression and enhanced disease resistance, while leaving early responses such as ROS burst and MAPK activation intact (Fig. 3a-e). These findings indicate that early PTI responses can proceed using preexisting pools of SMs, whereas late-stage amplification relies on de novo SM production.

The exclusive involvement of the TNL SADR1 in late PTI supports a two-tiered regulatory model for SM production during immune activation. SADR1 promotes PR1 expression and confers enhanced disease resistance^28^ without affecting early PTI markers (Fig. 3f,g), highlighting the specificity of TNL-mediated signaling branches. Together, these findings underscore the essential role of SOBIR1 in BAK1 recruitment to LRR-RPs, establish the requirement of SM binding to EDS1–PAD4 for robust PTI signaling, and propose a two-phase model of SM regulation— constitutive stores supporting early responses and inducible biosynthesis mediating late amplification—in plant innate immunity (Fig. 3h).

An important question is why *eds1* and *pad4* mutants defective in SM binding exhibit diminished PTI activation. Several explanations are possible, including reductions in basal transcript or protein abundance of key PTI components, disruption of signaling events required for early or late PTI responses, or loss of potential scaffold functions in which EDS1–PAD4 complexes organize or stabilize immune signaling assemblies. Because lowered basal salicylic acid (SA) can reduce expression of several LRR-RP pathway components^35^, we examined whether altered SA homeostasis contributes to the PTI defects of nlp20-impaired mutants. Total and free SA levels in unchallenged *eds1-2, pad4-1, adr1 triple*, and *pbl30 pbl31 pbl32* plants were comparable to those in Col-0 (Supplementary Fig. 8a,b), consistent with previous reports from mock treated genotypes^9,36-38^. Similarly, endogenous EDS1 and SOBIR1 protein abundances were unaltered in *sobir1-12* and *eds1-2* backgrounds (Supplementary Fig. 8c,d). Thus, compromised PTI activation in these mutants cannot be attributed to altered SA levels or baseline protein accumulation, consistent with earlier findings that transcripts and protein levels of immune-related genes are largely unchanged in unchallenged Col-0 and *pad4* plants^3^. Therefore, a key challenge for future work will be to determine whether EDS1–PAD4–ADR1 complexes function primarily as active signaling centers, organizational scaffolds, or multifunctional hubs orchestrating diverse layers of immune activation.

## Methods

### Plant materials and growth conditions

All mutants used in this study are in Arabidopsis accession Col-0 and all the lines are listed in Supplementary Table 1. *A. thaliana* plants were grown in soil in a growth chamber at 22ºC under short-day conditions of 8 h of light/16 h of dark.

### Peptides

Synthetic peptides (GenScript) were prepared as 10 mM stock solutions in 100% dimethyl sulfoxide (DMSO), and diluted in water to the desired concentration prior to use.

### Reactive oxygen species (ROS) burst assays

Leaves of 5-6-week-old Arabidopsis plants were cut into pieces (about 0.5x0.5 cm) and floated on water overnight. One leaf piece per well was transferred to a 96-well plates containing 20 µM luminol derivative L-012 (Wako Pure Chemical Industries Ltd, Osaka, Japan) and 20 ng ml-1 horseradish peroxidase (Applichem). Luminescence after treatment with peptides or water (as control) was measured with a luminometer (Mithras LB 940, Berthold) over 1 h in 2 min intervals. Total relative light unit production was determined by calculating the area under the scatter curve for the time points indicated.

### Measurement of ethylene production

Leaves of 6-7-week-old Arabidopsis plants were cut into pieces (about 0.5x0.5 cm) and floated on water overnight. Three leaf pieces were incubated in a sealed 6.5 ml glass tube with 0.4 ml 20 mM MES buffer, pH 5.7 and the indicated elicitors. Ethylene production was measured by gas chromatographic analysis (GC-14A; Shimadzu) of 1 ml air from the closed tube after incubation for 4 h.

### MAPK activation

Leaves of 5-6-week-old Arabidopsis plants were infiltrated with water or 1 µM nlp20. At the indicated time point, leaves were harvested and frozen in liquid nitrogen. MAPK activation assay was performed by immunoblotting with an anti-phospho p44/42 MAP kinase antibody (Cell Signaling Technology, 1:3,000 dilution).

### Quantitative reverse transcription PCR

Infiltrated leaves of 5-6-week-old Arabidopsis plants were harvested after 24 h for total RNA isolation with NucleoSpin® RNA Plus Kit (Macherey-Nagel). First strand cDNA was synthesized with RevertAid™ M-MuLV reverse transcriptase (Thermo Scientific) and qRT-PCR was performed with the iQ5 Multi-color real-time PCR detection system (Bio-Rad) using the SYBR Green Fluorescein Mix (Thermo Scientific) and gene-specific primers listed in Supplementary Table 2. Transcript levels of *PR1* were normalized to the transcript levels of the housekeeping gene *EF1α*.

### Pathogen infection assays

*Pseudomonas syringae* pv. *tomato* (*Pst*) DC3000 infection was performed as previously described^3^. Six-to 7-week-old plants were primed 24 h prior to infection by leaf infiltration of 1 µM peptide, or water (mock treatment). *Pst* DC3000 was resuspended in 10 mM MgCl_2_ at a density of 1x10^4^ cells ml-1 for Arabidopsis leaf infiltration. Bacterial growth was quantified 3 days after bacterial infiltration.

### Salicylic acid quantification

50 mg of leaves from 5–6-week-old *Arabidopsis* plants were ground in liquid nitrogen and sequentially extracted with 80% methanol containing phenyl-^13^C_6_ SA (internal standard, Sigma-Aldrich), followed by 20% methanol and 0.1% formic acid. Extractions were carried out at 10 °C with 10 minutes of shaking and centrifugation at 18,600 g. For free SA analysis, 5 μl of the combined extract was diluted 1:40 and analyzed by targeted LC-MS. For total SA, 100 μl of extract was hydrolyzed with concentrated formic acid at 99 °C for 1 hour, neutralized with 50% NaOH, and processed as above. Both free and total SA samples were analyzed using liquid chromatography–mass spectrometry (LC-MS), and the masses of SA and ^13^C_6_-SA were detected and quantified. Data analysis was performed using Sciex OS software (see detailed protocol in Supplementary File).

### NAM treatment

To assess the impact of NAM on PTI, Arabidopsis leaves were infiltrated with 50 mM NAM and cut into pieces after 30 min. After floating on water overnight, the leaf pieces were used for ROS burst and ethylene production assays as described above. Alternatively, leaves were co-infiltrated with 1 µM peptide with or without 50 mM NAM. After 15 min, the leaves were cut into pieces and directly transferred to the glass tube, then incubated for 5 h for ethylene measurement. For MAPK activation, *PR1* expression, and induced resistance assays, leaves were co-infiltrated with 1 µM nlp20 with or without 50 mM NAM and harvested at the indicated time points.

### Immunoprecipitation and protein blotting

Leaves of 5-6-week-old Arabidopsis plants expressing RLP23-GFP or RLP42-GFP were infiltrated with water or 1 µM nlp20 or pg13. After 5 min infiltration, the leaves were harvested and frozen in liquid nitrogen. Proteins were extracted at 1 mg ml-1 in extraction buffer [50 mM Tris-HCl, pH 7.4, 150 mM NaCl, 0.25% deoxycholic acid, 1% NP-40, 1 mM EDTA, proteinase inhibitor cocktail (Roche)]. Tagged proteins were immunoprecipitated for 1 h at 4°C using GFP-trap agarose beads (Proteintech) as previously described^3^. Immunoblots were performed using anti-GFP antibodies (Torrey Pines biolabs, 1:10,000) or anti-BAK1 antibodies (Agrisera, 1:10,000), followed by HRP conjugated anti-Rabbit (Agrisera, 1:20,000 ). Chemiluminescence was detected with the ECL Western blotting detection system (GE Healthcare) and a CCD camera (Amersham Imager 600).

Immunoblotting was performed to determine the background levels of proteins SOBIR1 (Agrisera, 1:1,000), EDS1 (Agrisera, 1:1,000), MPK3 (Sigma, 1:5,000), MPK4 (Sigma, 1:2,000), and MPK6 (Sigma, 1:1,000) using the same protein extracts for immunoprecipitation or MAPK activation assays. The intensities of examined proteins and corresponding loading by Ponceau S staining were measured using ImageJ.

## Data availability

All study data are included in the article and/or Supplementary file.

## Acknowledgements

We thank P. Jacob and J. L. Dangl for providing the *sadr1-c2* seeds. This work was supported by Deutsche Forschungsgemeinschaft (grants CRC1101-D10, Nu70/19-1, TRR356-B05 to T.N.). Metabolite analytics in ZMBP central facilities unit was funded by Deutsche Forschungsgemeinschaft (Project number 442641014).

## Author contributions

T.N. and L.Z. conceived and designed the experiments; J.F., D.J., C.H., E.v.R-L., F.L., and L.Z. conducted experiments; E.v.R-L., M.T., J.E.P., T.N., and L.Z. analyzed data; and T.N. and L.Z. wrote the manuscript. All authors discussed the results and commented on the manuscript.

## Additional information

Supplementary files include Supplementary Fig. 1-8 and Supplementary Table 1 and 2.

