## Supplemental files for "Small Molecule Binding to EDS1/PAD4 in LRR-RP-Mediated Pattern-Triggered Immunity"

**Supplementary files**

**Methods**

**Salicylic acid quantification.** 50 mg of leaves from 5–6-week-old *Arabidopsis* plants were harvested, ground in liquid nitrogen, and sequentially extracted with 200 µl 80% MeOH (with phenyl-$^{13}C_6$ salicylic acid standard, Sigma-Aldrich), 200 µl 20% MeOH, and 400 µl 0.1% formic acid (FA) in $H_2O$. Extractions were performed at 10 °C with 10 min shaking and centrifugation (18,600 g). For free SA analysis, 5 µl of the combined extract was diluted 1:40 and analyzed by targeted LC-MS. For total SA, 100 µl of the extract was hydrolyzed with 8 µl concentrated FA at 99 °C for 1 h, neutralized with 6 µl 50% NaOH, and processed as above. The LCMS profiling analysis was performed using a Micro-LC M5 (Trap and Elute) and a QTRAP6500+ (Sciex) operated in MRM mode (MRM transitions: SA (1) quantifier ion (m/z) Q1/Q3 137/93, collision energy CE -20V; (2) (m/z) Q1/Q3 137/65, CE -40V; $^{13}C_6$ Sa (1) quantifier ion (m/z) Q1/Q3 143/99, CE -20 V; (2) (m/z) Q1/Q3 143/70), CE -20 V). Declustering potential (DP) was set to –40 V for all transitions, with a dwell time of 5 ms for each MRM. Chromatographic separation was achieved on a Luna Omega Polar C18 column (3 µm; 100 Å; 150×0.5 mm; Phenomenex) and a Luna C18(2) trap column (5 µm; 100 Å; 20×0.5 mm; Phenomenex) with a column temperature of 58 °C and an injection volume of 50 µl.. The following binary gradient was applied for the main column at a flow rate of 14 µl min$^{-1}$: 0 - 0.2 min, isocratic 85% A; 0.2 – 4.5 min, linear from 85% A to 5% A; 4.5 - 5 min, isocratic 5% A; 5 - 6 min, linear from 5% A to 85% A; 6 - 7 min, isocratic 85% A (A: water, 0.1% aq. formic acid; B: acetonitrile, 0.1% aq. formic acid; all solvents were LC-MS grade). The samples were concentrated on the trap column using the following conditions: flow rate 25 µl min$^{-1}$: 0 - 2.5 min isocratic 95% A; 2.5 min start main gradient; 2.5 - 2.7 min isocratic 95% A. Analytes were ionized using an Optiflow Turbo V ion source equipped with a SteadySpray T micro electrode in positive (ion spray voltage: -4500 V) ion mode. Following additional instrument settings were applied: nebuliser and heater gas, nitrogen, 25 and 45 psi; curtain gas, nitrogen, 30 psi; collision gas, nitrogen, medium; source temperature, 200 °C; entrance potential, -10 V; collision cell exit potential, -10V. The Sa content in each sample was normalized against the $^{13}C_6$ Sa standard values. All data were integrated using the Sciex OS vendor software.

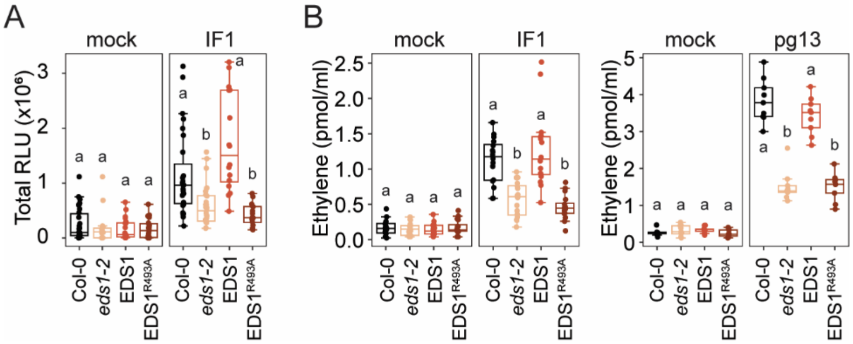


**Fig. S1.** SM binding to EDS1 is required for LRR-RP-mediated early PTI. (A) Total reactive oxygen species (ROS) production over 40 minutes in Col-0, *eds1-2*, and wild-type *EDS1* or the SM-binding mutant *EDS*$^{1R493A}$ complementation lines, following treatment with water (mock) or 1 μM IF1. Data represent three independent experiments. (B) Elicitor-induced ethylene production in Col-0, *eds1-2*, *EDS1*, and *EDS1*$^{R493A}$. For IF1 and pg13, n = 16 and 9 from four and three independent experiments, respectively. Letters indicate statistically significant differences among plant genotypes' responses to each elicitor, determined by the Kruskal-Wallis test followed by post-hoc Dunn's test (P < 0.01).

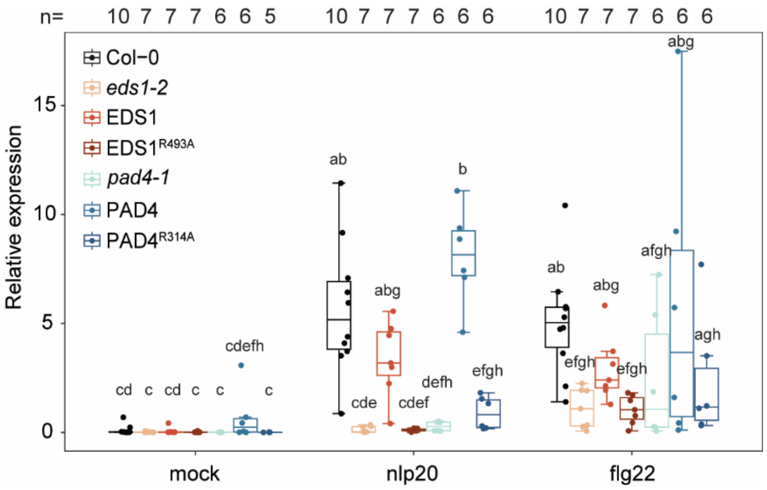


**Fig. S2.** Elicitor-induced PR1 expression. Leaves were treated with water (mock) or 1 µM nlp20 or flg22 and sampled 24 h after treatment. Relative expression of *PR1* was normalized to the levels of *EF-1α* transcript. Letters indicate statistically significant differences, determined by the Kruskal-Wallis test followed by post-hoc Dunn's test ($P < 0.05$).

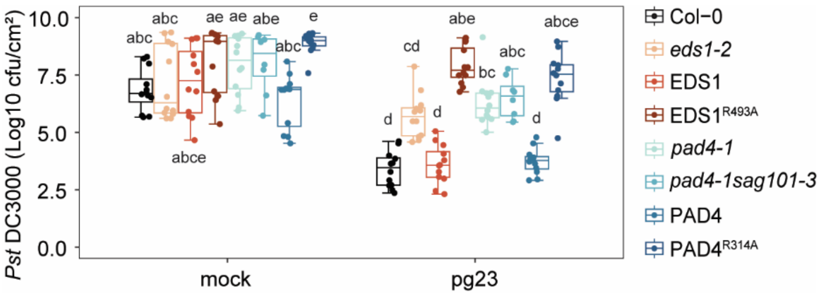


**Fig. S3.** Pg23-induced defense against *Pst* DC3000. Leaves were treated with water (mock) or 1 μM pg23 24 hours prior to *Pst*DC3000 infection. Bacterial titers were quantified in leaf extracts at 3 days post-inoculation (dpi). n = 12 biological replicates (each comprising two leaf discs). Letters indicate statistically significant differences, determined by the Kruskal-Wallis test followed by post-hoc Dunn's test ($P < 0.01$).

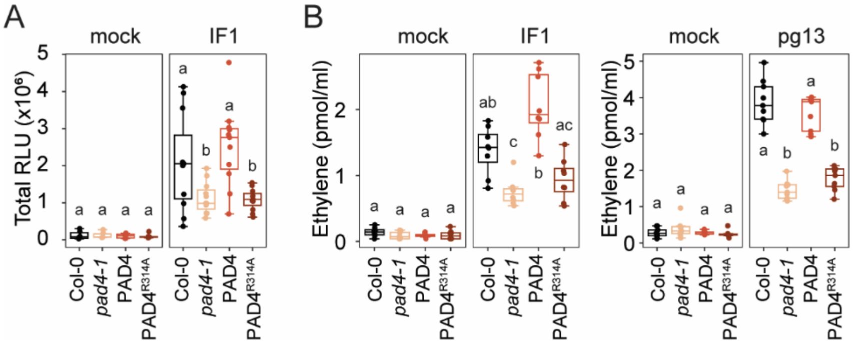


**Fig. S4.** SM binding to PAD4 is required for LRR-RP-mediated early PTI. (A) Total reactive oxygen species (ROS) production over 40 minutes in Col-0, *pad4-1*, and wild-type *EDS1* or the SM-binding mutant *EDS1*$^{R493A}$ complementation lines, following treatment with water (mock) or 1 μM IF1. Data represent three independent experiments. (B) Elicitor-induced ethylene production in Col-0, *eds1-2*, *EDS1*, and *EDS1*$^{R493A}$. For IF1 and pg13, n = 8 and 9 from two and three independent experiments, respectively. Letters indicate statistically significant differences among plant genotypes' responses to each elicitor, determined by the Kruskal-Wallis test followed by post-hoc Dunn’s test (A, $P < 0.05$; B, $P < 0.01$).

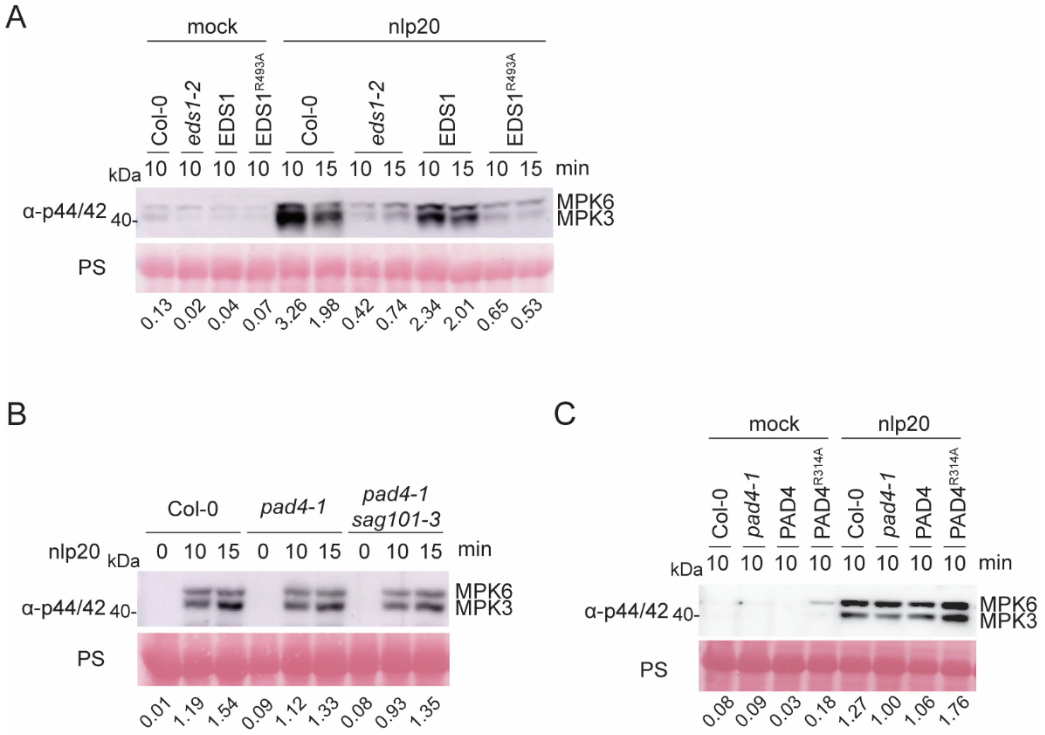


**Fig. S5.** EDS1, but not PAD4, is required for nlp20-triggered MAPK activation. Leaves from the indicated plant genotypes were treated with water (mock) or 1 µM nlp20, and MAPK activation was detected by immunoblot using an anti-p44/42 MAP kinase antibody. Ponceau S (PS) staining was used as a loading control. Relative band intensities of MPK6 and MPK3, normalized to loading (PS staining), are shown below the blots. Experiments were performed three times with consistent results.

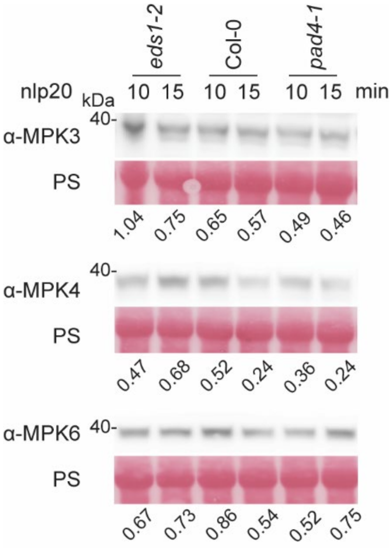


**Fig. S6.** Protein levels of MPK3, MPK4, and MPK6 in Col-0, *eds1-2*, and *pad4-1* mutant plants. Immunoblot analysis was performed using anti-MPK3, anti-MPK4, and anti-MPK6 antibodies on protein extracts from Arabidopsis leaves following 10- and 15-minute treatment with 1 μM nlp20. Ponceau S (PS) staining was used as a loading control. Relative band intensities of MPK3, MPK4, and MPK6, normalized to loading (PS staining), are shown below the blots.

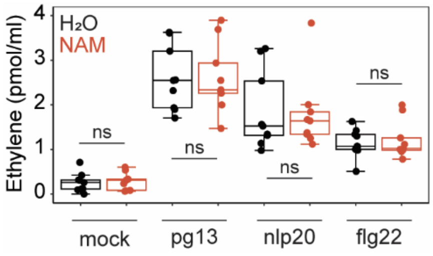


**Fig. S7.** NAM treatment prior to elicitor application does not affect early PTI responses. Ethylene accumulation in Col-0 pre-treated with water or 50 mM NAM 16 hours before mock or elicitor treatment (N = 9 from 3 independent experiments). No statistically significant difference (ns) by two-tailed Student’s t-test.

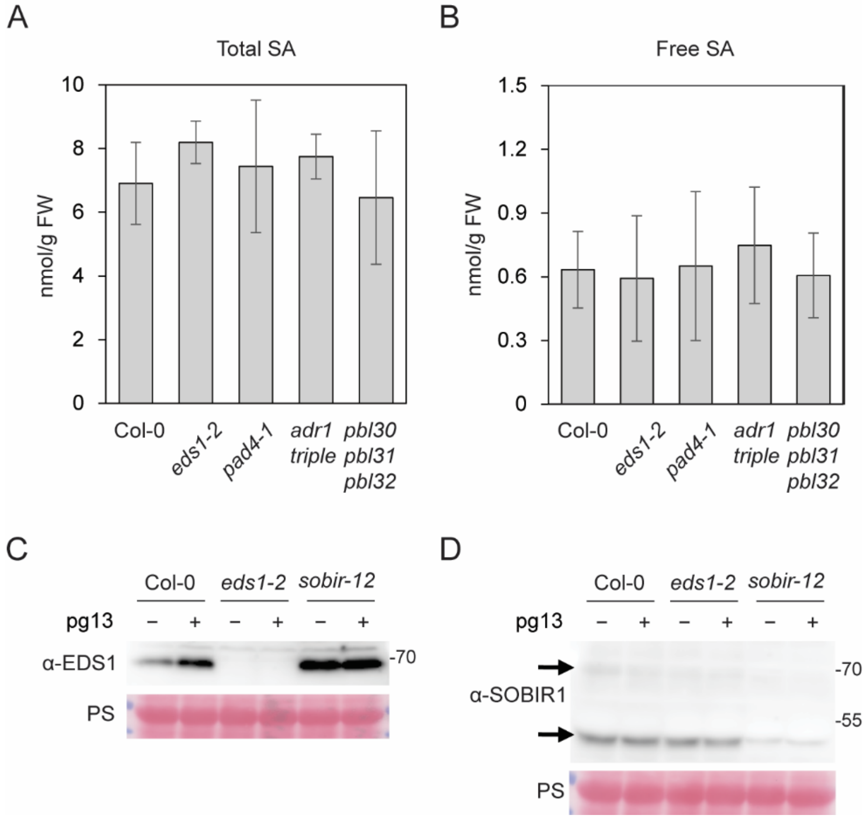


**Fig. S8.** Salicylic acid (SA) levels and protein expression in Arabidopsis. (A, B) Quantification of total SA (A) and free SA (B) in untreated 5-6-week-old leaves of Col-0 and mutant plants. Five individual plants per genotype were used for measurements. Bars represent means ± standard deviation. (C, D) Immunoblot analysis using anti-EDS1 (C) and anti-SOBIR1 (D) antibodies on protein extracts from Arabidopsis leaves treated for 5 minutes with water (−) or 1 μM pg13 (+). Ponceau S (PS) staining was used as a loading control.

**Table S1.** Arabidopsis lines used in this study.

| Line | Locus | Description | Reference |
|---|---|---|---|
| Col-0 p35S:RLP23-GFP | | overpressing RLP23-GFP in Col-0 | This study |
| Col-0 p35S:RLP42-GFP | | overpressing RLP42-GFP in Col-0 | This study |
| *eds1-2* p35S:RLP23-GFP | *At3g48090* | overpressing RLP23-GFP in *eds1-2* | This study |
| *eds1-2* p35S:RLP42-GFP | *At3g48090* | overpressing RLP42-GFP in *eds1-2* | This study |
| *pad4-1* p35S:RLP23-GF*P* | *At3g52430* | overpressing RLP23-GFP in *pad4-1* | This study |
| *pad4-1* p35S:RLP42-GF*P* | *At3g52430* | overpressing RLP42-GFP in *pad4-1* | This study |
| *sobir1-12* p35S:RLP23-GFP | *At2g31880* | overpressing RLP23-GFP in *sobir1-12* | This study |
| *sobir1-12* p35S:RLP42-GFP | *At2g31880* | overpressing RLP42-GFP in *sobir1-12* | This study |
| *eds1-2* | *At3g48090* | Polymorphism 1009135505, eds1-2, introgressed into Col-0 | Bartsch et al. (2006) |
| *eds1* cEDS1 | *At3g48090* | Col-0 *eds1-2* complemented with c*EDS1* (cloned from cDNA) | Bhandari et al. (2019) |
| *eds1* cEDS1$^{R493A}$ | *At3g48090* | Col-0 *eds1-2* complemented with cEDS1$^{R493A}$, harboring a mutation required for SM binding | Bhandari et al. (2019) |
| *pad4-1* | *At3g52430* | Polymorphism, 4770301, *pad4-1* | Jirage, et al. (1999) |
| *pad4-1 sag101-3* | *At3g52430, At5g14930* | *pad4-1 sag101-3* double knockout line | Cui, et al. (2018) |
| *pad4-1 sag101-3* gPAD4 | *At3g52430, At5g14930* | *pad4-1 sag101-3* complemented with gPAD4 (cloned from genomic DNA) | Dongus, et al. (2022) |
| *pad4-1 sag101-3* gPAD4$^{R314A}$ | *At3g52430, At5g14930* | *pad4-1 sag101-3* complemented with gPAD4 R314A, harboring a mutation required for SM binding | Dongus, et al. (2022) |
| *sadr1-c2* | *At4g36150* | loss-of-function mutant by CRISPR-Cas9 | Jacob, et al. (2023) |

**Table S2.** qRT-PCR primers used in this study.

| Primer name | Sequence |
|---|---|
| qPR1_F | CGCTGCGAACACGTGCAATG |
| qPR1_R | CCACGAGGATCATAGTTGCAAC |
| qEF1a_F | GAGGCAGACTGTTGCAGTCG |
| qEF1a_R | TCACTTCGCACCCTTCTTGA |
